# CRISPR Cas13-based tools to track and manipulate endogenous telomeric repeat-containing RNAs in living cells

**DOI:** 10.1101/2021.12.03.471109

**Authors:** Meng Xu, Tafadzwa Chigumira, Ziheng Chen, Jason Tones, Rongwei Zhao, Kris Noel Dahl, David M. Chenoweth, Huaiying Zhang

## Abstract

TERRA, TElomeric Repeat-containing RNA, is a long non-coding RNA transcribed from telomeres. Emerging evidence indicates that TERRA regulates telomere maintenance and chromosome end protection in normal and cancerous cells. However, the mechanism of how TERRA contributes to telomere functions is still unclear, partially owing to the shortage of approaches to track and manipulate endogenous TERRA molecules in live cells. Here, we developed a method to visualize TERRA in live cells via a combination of CRISPR Cas13 RNA labeling and Suntag technology. Single-particle tracking reveals that TERRA foci undergo anomalous diffusion in a manner that depends on the timescale and telomeric localization. Furthermore, we used a chemically-induced protein dimerization system to manipulate TERRA subcellular localization in live cells. Overall, our approaches to monitor and control TERRA locations in live cells provide powerful tools to better understand its roles in telomere maintenance and genomic integrity.

## Introduction

Telomeres, the repetitive DNA sequences at chromosome ends, are coated by the Shelterin protein complex to protect them from incorrect fusion and recombination as DNA double-strand breaks (Maciejowski & De Lange, 2017; O’Sullivan & Karlseder, 2010; Palm & de Lange, 2008). In addition to Shelterin, TElomeric Repeat-containing RNAs (TERRAs) also play important roles in telomere integrity. TERRAs are transcribed from the subtelomeric regions towards chromosome ends by RNA polymerase II and are highly heterogeneous transcripts with sizes ranging from 100 nt to 9 kb in mammalian cells (Azzalin et al., 2007; Bettin et al., 2019; Schoeftner & Blasco, 2008). A growing body of studies indicates that TERRA actively regulates telomere function and maintenance (Azzalin et al., 2007; Chu et al., 2017; Deng et al., 2009; Feretzaki et al., 2020; Wang et al., 2015). Of note, TERRA has a multifaceted role for telomere maintenance, including facilitating telomere replication (Beishline et al., 2017; Petti et al., 2019; Silva et al., 2021) and heterochromatin formation at telomeres (Deng et al., 2009; Montero et al., 2018).

TERRA is also involved in telomere maintenance of cancer cells (Bettin et al., 2019; Chu et al., 2017; De Silanes et al., 2014). Actively maintaining telomere length is required for cancer cells to counteract the replicative barrier induced by telomere shortening in cell division for their immortality (Bonnell et al., 2021; Hanahan & Weinberg, 2011). While most human cancers acquire unlimited replication via reactivating the reverse transcriptase telomerase, 10%-15% of cancers use mechanism so-called alternative lengthening of telomeres (ALT) pathway where cancer cells hijack the homologous recombination-based DNA repair to extend telomeres (Claude & Decottignies, 2020; Dilley & Greenberg, 2015; Recagni et al., 2020). TERRA contributes to telomere maintenance in both types of cancer cells. In telomerase-positive cancer cells, TERRA directly regulates telomerase activity (Azzalin & Lingner, 2015; Lalonde & Chartrand, 2020). In ALT cancer cells, TERRA is uniquely upregulated and forms R-loops to promote the ALT telomere maintenance (Arora & Azzalin, 2015; Yeager et al., 1999; J. M. Zhang et al., 2019).

Although the critical role of TERRA for telomere integrity is well-established, the mechanism of how TERRA acts at the subcellular level is still unclear. In this regard, a better understanding of the spatiotemporal dynamics of TERRA can help define the functions of TERRA. So far, two methods have been reported to monitor endogenous TERRA in live cells. The first is integrating MS2 repeats into a telomere that transcribes TERRA so the transcribed TERRA can be visualized by a fluorescently tagged MS2 binding protein (Avogaro et al., 2018). However, this method can only be used to image TERRA transcribed from the engineered telomere, as opposed to all TERRA since TERRA is transcribed from multiple telomeres. Another live-cell method uses an engineered TERRA binding protein called mPUMt, the mutant of the Pumilio homology domain. (Yamada et al., 2016). By fusing mPUMt to split GFPs and imaging with a special microscopy technique called total internal reflection fluorescence microscopy, TERRA interaction with telomeres at the single-molecule level can be monitored. However, imaging with common microscopy techniques, such as epifluorescence and confocal microscopy, is not feasible with this method. Thus, better tools to track TERRA localization and dynamics in live cells are imperative.

In addition, TERRA also binds to other regions of the chromosome, and only a fraction of TERRA localizes to telomeres (Biffi et al., 2012; Chu et al., 2017; Diman & Decottignies, 2018; Mei et al., 2021; Yang et al., 2019). TERRA telomeric localization is tightly regulated: too much or too little results in telomere dysfunction, based on results obtained by manipulating TERRA-interacting proteins (De Silanes et al., 2010). Since those TERRA binding proteins are known to affect telomere integrity (Petti et al., 2019; Porreca et al., 2020), it is therefore not clear the direct contribution of telomere-bound TERRA in telomere function. Therefore, tools to manipulate TERRA localization are desirable for assessing the functional importance of TERRA localization.

Here, we developed a system to visualize endogenous TERRA in live cells based on CRISPR-dPspCas13b technology. Furthermore, to increase imaging efficiency, we amplified the signals via combination with the repeating peptide array (SunTag). Importantly, relying on this system, we monitored the dynamics of TERRA foci with single-particle tracking. Lastly, we combine the dCas13b-SunTag tool with a chemically induced protein dimerization system to control TERRA localization on telomeres.

## Results

### Design of guide RNA to image TERRA with CRISPR-cas13

To probe the endogenous TERRA, we utilized RNA-guided catalytically inactive Cas13b system (Figure 1A), which was reported to detect RNA in live cells (Yang et al., 2019). Given that TERRA is variable with tandem (UUAGGG)_n_ repeats sequence, there is no sequence specificity for guide RNA recognition. Therefore, we designed three guide RNAs with different lengths ranging from 22 to 30 nucleotides (nt) (Figure 1B). With the addition of guide RNA, EGFP-fused dPspCas13b indeed forms visible foci in the nucleoplasm in addition to obvious nucleolar signals in live cells (Figure 1B). Significantly, the length of guide RNA determines the RNA-labeling efficiency. We found that the shortest guide RNA with 22 nt induces more visible foci than the longer ones (Figure 1C). To verify that those are TERRA foci, we employed RNA fluorescent in situ hybridization (FISH) with TERRA probe in fixed cells (Figure 1D). As expected, the dPspCas13b foci are all labeled by the TERRA FISH probe. In addition, the TERRA signal is decreased after treatment with Ribonuclease, indicating the RNA-binding specificity of the TERRA probe. This suggests that the CRISPR-dCas13 system labels TERRA properly.

**Figure 1.**
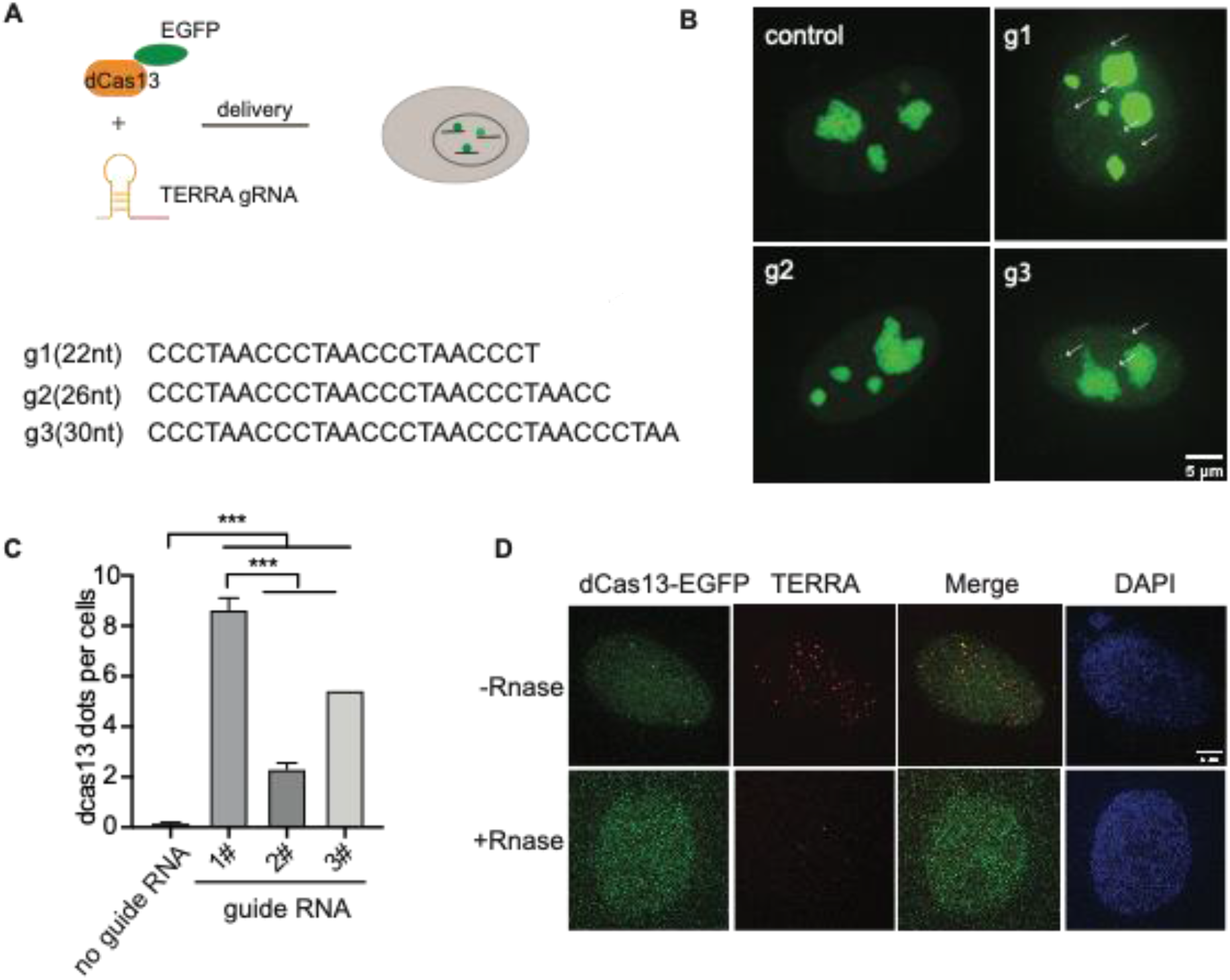
CRISPR-dCas13 enables visualization of TERRA in live cells. (A) Overview of CRISPR-dCas13-mediated TERRA labeling. (B) Representative images of dCas13b-EGFP with three different guide RNAs for TERRA and guide for non-target control RNA. Arrows indicate dCas13b-EGFP labeled TERRA. (C) Quantification of total foci per cell indicated by dCas13 with different guide RNAs (mean ± SEM, unpaired t-test). N≥100 for each group. ***, p<0.001. (D) RNA FISH shows colocalization of dCas13b-EGFP with TERRA foci (red). DAPI detects nuclear DNA. Arrow indicates colocalization of TERRA FISH and dCas13b-EGFP.

### Increase labeling efficiency with SunTag

Although EGFP-fused dCas13b detects TERRA, the signal is weak compared to the non-specific signals in nucleoli, restricting its utility for dynamic imaging of TERRA in live cells. We expect that those nucleolar signals are caused y protein aggregation in nucleoli. To improve TERRA imaging efficiency, we combined the SunTag technology with the CRISPR-dcas13 system to amplify the TERRA signal. The synthetic SunTag scaffold, including five tandem GCN4, is fused to dCas13b to recruit up to five GFP copies via scFV (Figure 2A). Additionally, we replaced EGFP with sfGFP, a form of superfolder GFP, to increase its solubility (Pédelacq et al., 2006). As visualized in Figure 2B with FISH, those visible foci indicated by dCas13b-SunTag are all TERRA positive as well (Figure 2B). Notably, the non-specific fluorescent signal in nucleoli is largely decreased with the dCas13-SunTag system. Significantly, in contrast to the original dCas13b strategy, the combination with SunTag largely increases the TERRA detection rate from around 5% to 38% (Figure 2C). Also, TERRA foci detected by dCas13b-SunTag-sfGFP are bigger and brighter than dCas13b-EGFP dots, owing to five GFP copies binding to SunTag via scFV. To demonstrate that the dCas13b signal is indeed from TERRA RNA, we analyzed the percentage of dCas13 proteins detected by the TERRA probe. The data shows that 77% of dCas13-GFP and 89% of dCas13-SunTag-sfGFP are TERRA positive (Figure 2D). The results indicate that the SunTag technology with sfGFP improves TERRA labeling efficiency.

**Figure 2.**
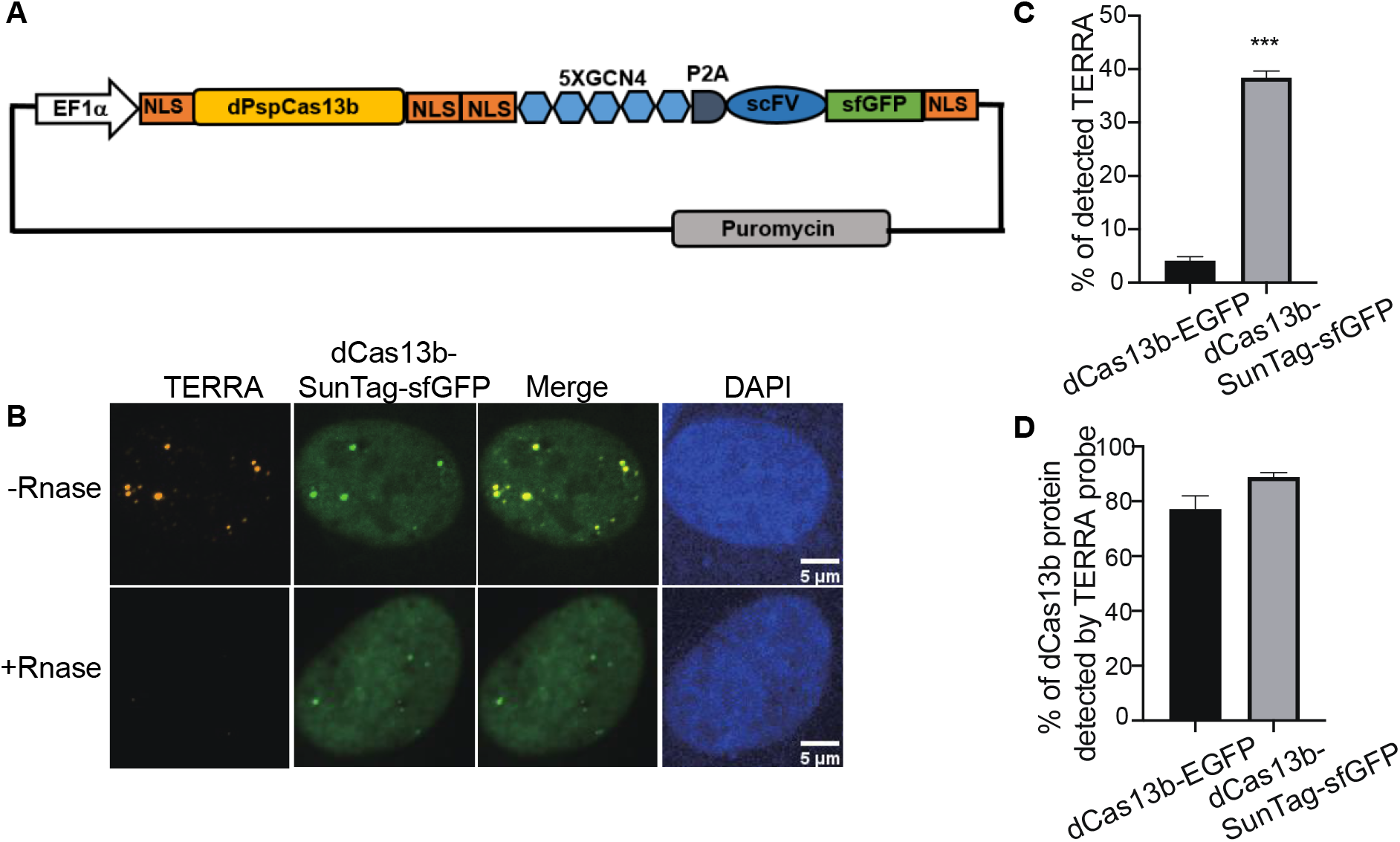
SunTag increases TERRA labeling efficiency. (A) Plasmid map of dCas13b tagged with SunTag, including five repeats of GCN4, single-chain variable fragment antibody (scFV) followed by superfolded GFP (sfGFP). (B) TERRA RNA FISH shows co-localization of dCas13b-SunTag-sfGFP with TERRA probe (red). (C) Quantification for percentage of TERRA FISH foci labeled by dCas13b-EGFP and dCas13b-SunTag-sfGFP (mean ± SEM, unpaired t-test). N≥80 for each group. ***, p<0.001. (D) Quantification for percentage of dCas13b-EGFP and dCas13b-SunTag-sfGFP labeled by TERRA FISH (mean ± SEM). N≥30 for each group.

### Timescale and location-dependent TERRA foci movement revealed with single-particle tracking

The capacity of dCas13b-SunTag-sfGFP to detect TERRA foci in live cells prompted us to monitor TERRA foci movement with single-particle tracking. The movement of many structures in the human cell nucleus, such as nanoparticles, PML nuclear bodies, and telomeres, are shown to depend on timescale, owning to particle confinement within the chromatin cages at small timescales and particle hopping between cages at large timescales (Tseng et al., 2004, Jegou et al., 2009). To determine whether TERRA foci movement differs with timescales, we generated TERRA foci trajectories at two timescales: 0-1 second and 10-100 seconds (Figure 3A). Furthermore, since a subset of TERRA foci co-localizes with telomeres (Biffi et al., 2012; Mei et al., 2021; Yang et al., 2019), we aimed to determine whether telomeric localization affects TERRA foci dynamics by imaging telomeres through mCherry fused to Shelterin component TRF1 while tracking TERRA foci (Figure 3B).

**Figure 3.**
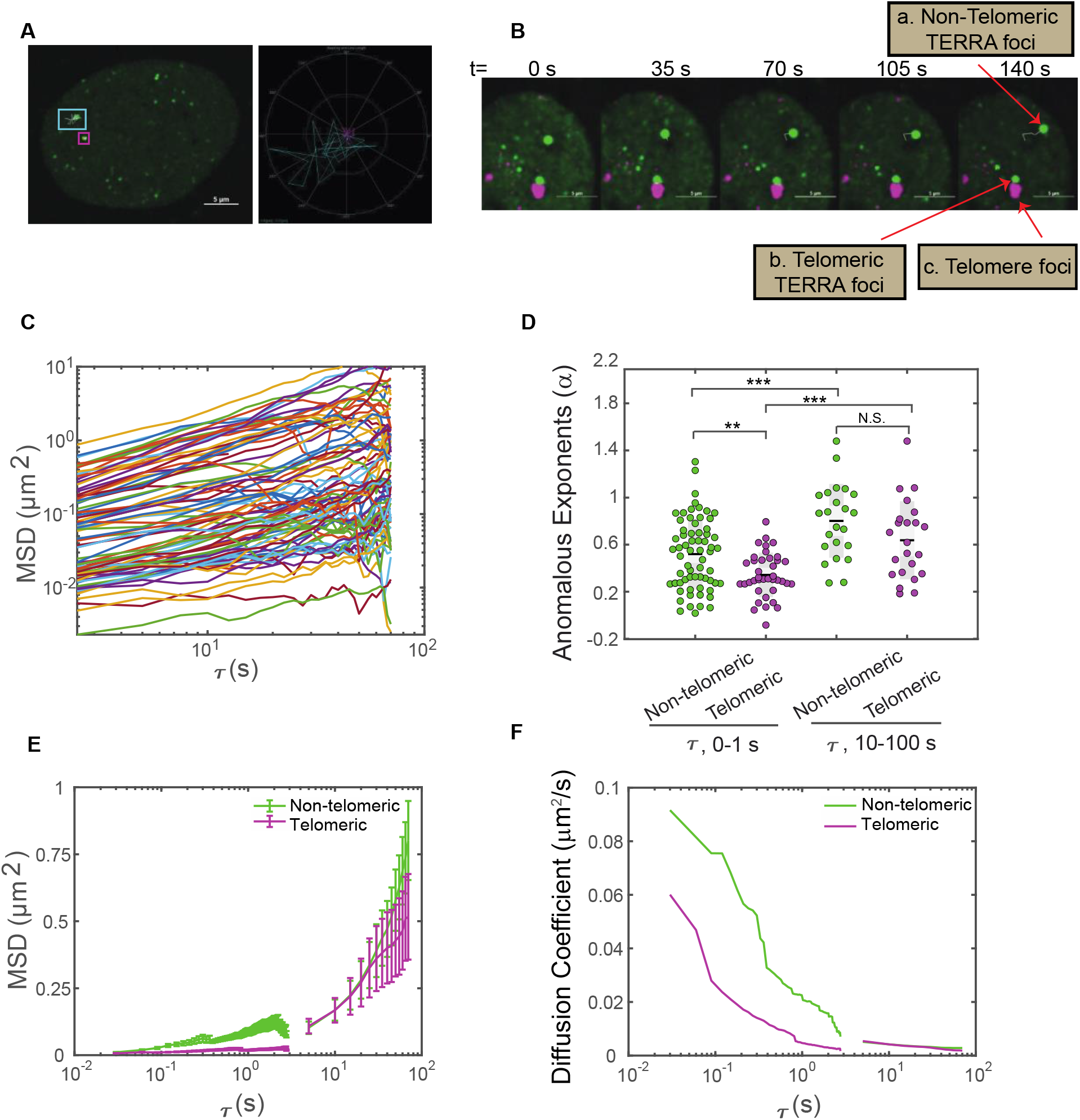
Single-particle tracking of TERRA foci mobility in live cells. (A)Left, U2OS cells expressing dCas13b-SunTag-sfGFP with an overlay of representative TERRA foci tracks. Right, trajectories of the TERRA foci highlighting the areas explored by the two selected TERRA foci in the left image. (B) Time-series images showing a representative TERRA focus (a) that is not localized to telomeres (called non-telomeric) and a TERRA focus (b) whose motions are in tandem with the neighboring telomere foci (c) and so is taken to be colocalized with the telomere (called telomeric). (C) Representative Mean Square Displacement (MSD) trajectories plotted versus lag time (τ) on log-log coordinates for TERRA foci. Each line represents one TERRA track. (D) The anomalous exponents evaluated from the MSD plots on two timescales 0-1s and 10-100s for telomeric and non-telomeric TERRA foci, each dot represents one TERRA track. *, p<0.05, **, p<0.01, and ***, p<0.001. (E) MSDs averaged at each *τ* for telomeric and non-telomeric TERRA foci. (F) Time-dependent diffusion coefficients evaluated from the average MSD plots for telomeric and non-telomeric TERRA foci over lag time *τ*.

Overall, TERRA foci move heterogeneously, with some diffusing within a small area while others explore a region several times larger (Figure 3A). To quantify whether and how the heterogeneity of TERRA foci movement depends on the timescale and telomeric localization, we generated the mean-squared displacement (MSD) curves over the lag time *τ* for telomeric and non-telomeric TERRA foci at the two timescales (Figure 3C, Figure S1A, B). To assess the mode of motion, i.e., whether it deviates from normal diffusion as seen for other molecules/structures in the nucleus (Woringer & Darzacq, 2018), we fitted the MSD curves of TERRA foci to the equation for anomalous diffusion, MSD=Kτ^α^, where K is the generalized diffusion coefficient, *τ* is the lag time, and α is the anomalous exponent (α =1 for normal diffusion, α <1 for anomalous diffusion, and α >1 for active diffusion). Average α for telomeric and non-telomeric TERRA foci at the 0-1 s and 10-100 s timescales are 0.34, 0.52, 0.64, and 0.7, respectively, indicative of overall anomalous diffusion (Figure 3D, Table S1). Lower α for telomeric and non-telomeric TERRA foci at the 0-1 s timescale than their counterparts at the 10-100 s timescale agrees with reported caging of particles in the chromatin network at small timescales (Tseng et al., 2004). Lower α of telomeric TERRA foci than non-telomeric foci at the 0-1s timescale, but not at the 10-100s timescale. This suggests that the local telomere environment confines TERRA foci more than other regions at small timescales, but attachment to telomeres does not alter TERRA foci hopping between different chromatin cages at large timescales.

To compare the difference in mobilities of TERRA foci with different anomalous exponents, we calculated the mean MSD at each *τ* and converted it to a time-dependent diffusion coefficient, D, following MSD=4D*τ* (Figure 3EF). Interestingly, TERRA foci diffusion coefficients decay quickly in the timescale of 0-1s but seem to plateau at 10-100s, similar to the behavior of other structures in the nucleus (Tseng et al., 2004). In addition, at the 0-1 s timescale, diffusion coefficients of telomeric TERRA foci are smaller than non-telomeric TERRA foci (mean 0.013 vs. 0.037 µm^2^/s), which suggests that the local telomere environment not only makes TERRA foci move more anomalously but also slower. However, at the 10-100 s timescale, no significant difference in diffusion coefficients for telomeric and non-telomeric foci is observed (mean 0.0033 vs. 0.0029 µm^2^/s, Figure S1, Table S1), indicating attachment to telomeres affects neither the mode nor magnitude of TERRA movement. Taken together, the dCas13b-SunTag-sfGFP system enabled us to monitor TERRA foci movement with single-particle tracking and revealed its dependence on timescale and telomeric localization.

### Control TERRA telomeric localization with chemical dimerization tools

In addition to monitoring TERRA localization and motion, we exploit our dCas13b-SunTag tool to control TERRA localization by combining it with a small molecule-mediated protein dimerization system we developed (Ballister et al., 2014; H. Zhang et al., 2017, 2020). This system is based on two linked ligands, TMP (Trimethylolpropane) and Halo, that can interact with the protein eDHFR and Haloenzyme, respectively (Figure 4A). We fused Haloenzyme to the telomere binding protein TRF1 to localize the dimerizers to telomeres and eDHFR to SunTag. Adding the chemical dimerizer, TMP-NVOC (6-nitroveratryl oxycarbonyl)-Halo, would recruit dCas13b protein, and thus TERRA, to telomeres (Figure 4A). By using an antibody against telomere binding protein TRF2 to label telomeres and TERRA FISH to confirm TERRA localization, we observe a basal level of TERRA localization on telomere without dimerizers, consistent with other studies (Figure 4B)(Azzalin et al., 2007; Chu et al., 2017). After adding dimerizers, the colocalization of TERRA on telomere increased two-fold (Figure 4C), whose effect on telomere function awaits to be determined. Meanwhile, the colocalization of dCas13b proteins on telomere is up to 80% from 6% with the dimerizer (Figure 4D), suggesting the dimerization efficiency is high. Overall, the dCas13b-SunTag is compatible with the protein dimerization system for spatiotemporal enrichment of TERRA on telomere for functional studies.

**Figure 4.**
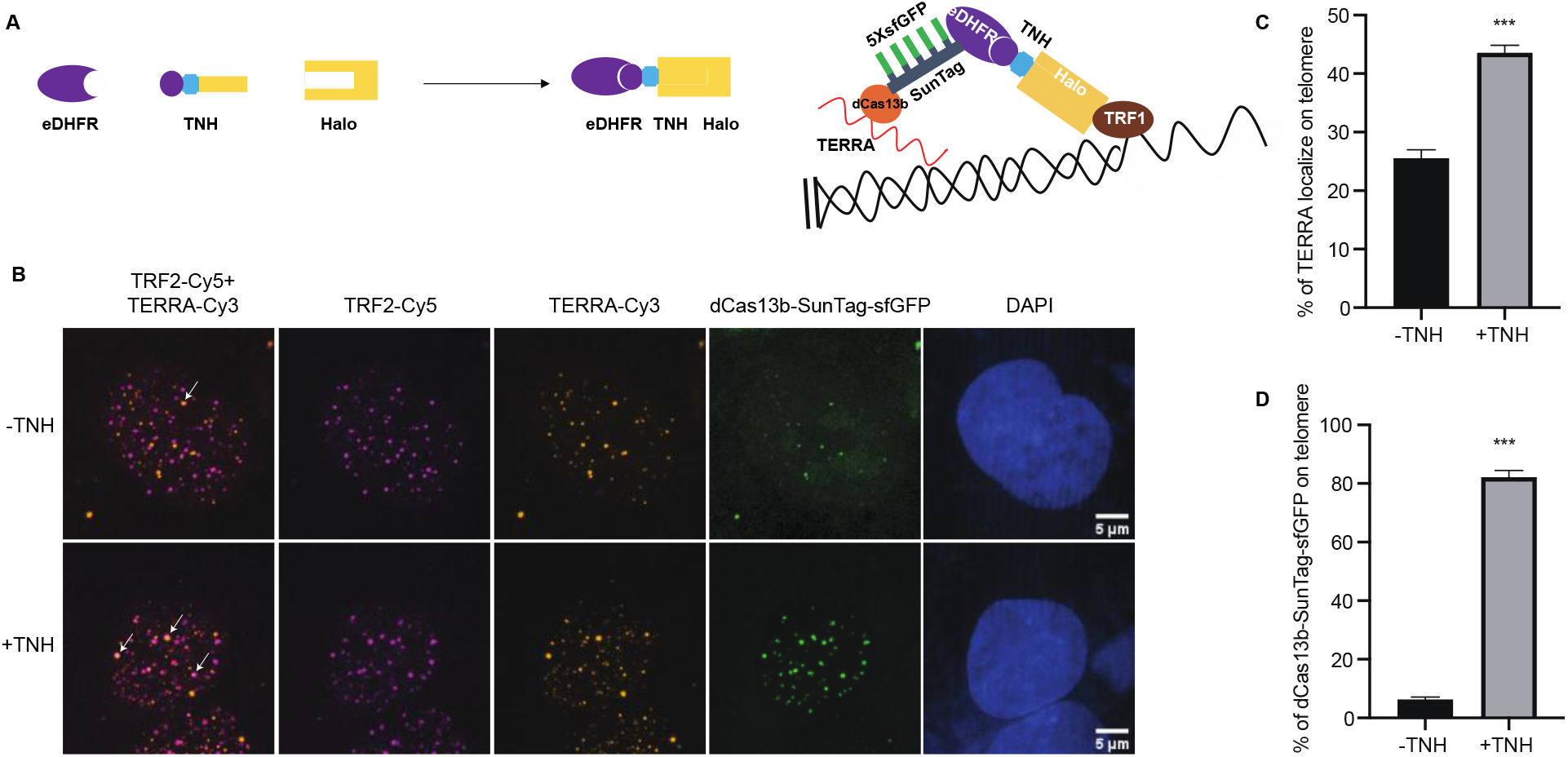
Protein dimerization system recruited TERRA to telomeres through Cas13b. (A) Dimerization schematic: SunTag-sfGFP is fused to eDHFR, and TRF1 is fused to Halo. The dimerizer is TNH: TMP-NVOC (6-nitroveratryl oxycarbonyl)-Halo (left panel), with TMP binds eDHFR, and Halo ligand binds to haloenzyme. Schematic diagram of recruitment of TERRA on telomere via dimerization system (right panel). (B) Representative images of TERRA localization on telomere verified by TERRA FISH. Arrow indicates colocalization of TERRA-Cy3 and TRF2-Cy5. (C) Quantification of the percentage of TERRA localized on telomere (mean ± SEM, unpaired t-test). N≥60 for each group. ***, p<0.001. N.S., not significant. (D) Quantification of the percentage of dCas13b-SunTag-sfGFP localized on telomere indicated by TRF2-Cy5 (mean ± SEM, unpaired t-test). N≥30 for each group. ***, p<0.001.

## Discussion

There is a growing consensus that characteristic distributions and dynamics of TERRA correlate with its function. Indeed, apart from the transient localization of TERRA on telomeres, TERRA molecules have been reported to bind chromatin throughout the genome (Chu et al., 2017; Marión et al., 2019). Thus, robust tools to track and manipulate the spatiotemporal dynamics of TERRA are vital to understanding TERRA functions. In this study, we developed a live-cell imaging method to visualize endogenous TERRA localization and dynamics by using CRISPR-Cas13 techniques combined with the SunTag system. In addition, we successfully integrated the Cas13 TERRA labeling with a protein dimerization system to control TERRA localization.

Compared with currently published tools, our methods have a few advantages. First, compared to MS2 integrated into one telomere, dCas13b can detect endogenous TERRA molecules universally (Avogaro et al., 2018). Secondly, compared with the assay based on the TERRA-recognizing domain mPUMt that relies on a highly specialized TIRF microscope system which restricts its broad utility (Yamada et al., 2016), the dCas13b-SunTag system can visualize TERRA with regular confocal microscope easily. Thirdly, the SunTag system vastly amplified the TERRA signal (Tanenbaum et al., 2014), offering better photobleaching resistance to enable long-term imaging of TERRA in live cells, such as the single-particle tracking of TERRA foci demonstrated here.

The timescale and location dependence of TERRA foci movement suggest that TERRA foci mobility can be used to reflect its local physical-chemical environment to provide insights into its biological functions. Finally, the dCas13b-SunTag system enabled us to manipulate TERRA localization with our chemical dimerization system, which can be used to dissect location-specific TERRA functions.

Since long non-coding RNAs are particularly effective at nucleating condensates by interacting with RNA-binding proteins in the nucleus (Frank & Rippe, 2020; Jain & Vale, 2017; H. Zhang et al., 2015), it is possible that TERRA phase separates with TERRA binding proteins to form condensates for telomere maintenance in normal or cancerous cancer cells. In addition, recent studies reported phase separation plays an essential role in telomere elongation in the ALT cancer cells (Min et al., 2019; H. Zhang et al., 2020). Telomere binding protein TRF2, the critical protein in protecting telomere integrity, was also shown to phase separate with TERRA (Soranno et al., 2021). The tools developed here to monitor and manipulate TERRA localization in live cells can be readily used to assess TERRA phase behavior and its functional significance.

## Materials and Methods

### Plasmids

The plasmids of expression dPspCas13b, guide RNA and GCN4-scFv-sfGFP (SunTag) were all purchased from addgene (#132397, #103854, #60906). To construct the dCas13b-SunTag-sfGFP, GCN4-scFv-sfGFP were amplified from addgene plasmid #60906 and introduced into plasmid#132397 through in-fusion cloning (Takara Bio). All other plasmids in this study are derived from a plasmid that contains a CAG promoter for constitutive expression, obtained from E. V. Makeyev (Khandelia et al., 2011).

### Cell culture

All experiments were performed with U2OS acceptor cells, originally obtained from E.V. Makayev (Khandelia et al., 2011). Cells were cultured in growth medium (Dulbecco’s Modified Eagle’s medium with 10% FBS and 1% penicillin–streptomycin) at 37 °C in a humidified atmosphere with 5% CO2. The constructs used in this manuscript, including mCherry-TRF1, dCas13-EGFP, dCas13-SunTag-sfGFP and Halo-TRF1, were transiently expressed by transfection with Lipofectamine 3000 (Invitrogen) 24 hours prior to imaging, following the manufacturer’s protocol.

### TERRA fluorescence in situ hybridization (FISH)

TERRA FISH assay was performed as previously described (Flynn et al., 2011). Briefly, cells were washed twice with cold PBS and treated with cytobuffer (100 mM NaCl, 300 mM sucrose, 3 mM MgCl2, 10 mM PIPES pH 7, 0.1% Triton X-100, 10 mM vanadyl ribonucleoside complex) for 7 min on ice. Cells were rinsed with cytobuffer (100 mM NaCl, 300 mM sucrose, 3 mM MgCl2, 10 mM PIPES pH 7, 0.1% Triton X-100, 10 mM vanadyl ribonucleoside complex) for 7 min at 4 °C, fixed in 4% formaldehyde for 10 min at room temperature, followed by permeabilization in 0.5% Triton X-100 for 10 min. For the control group, cells were then digested with RNaseA 200 mg/ml in PBS for 30 min at 37°C and were washed twice with PBS for 5 min. After incubation with blocking solution containing 1% BSA for 1 h, cells were then dehydrated in a series of ethanol washes 70%, 85%, 100% for 5 min each at room temperature, and the coverslips were dried at room temperature. 20 nM Telo Miniprobe SCy3 short probe in hybridization buffer (50% formamide, 2x SSC, 2mg/ml BSA, 10% dextran sulfate) was added to coverslips and then placed in a humidified chamber at 39°C overnight. The following day, the coverslips were washed in 2x SSC +50% formamide three times at 39°C for 5 min each, three times in 2xSSC at 39°C for 5 min each, and finally one time in 2x SSC at room temperature for 10 min. The coverslips were mounted on glass microscope slides with Vectashield mounting medium containing DAPI and analyzed with microscopy.

### Image acquisition

For live imaging, cells were seeded on 22×22mm glass coverslips (no. 1.5; Fisher Scientific) coated with poly-D-lysine (Sigma-Aldrich) in single wells of a 6-well plate. When ready for imaging, coverslips were mounted in magnetic chambers (Chamlide CM-S22-1, LCI) with cells maintained in L-15 medium without phenol red (Invitrogen) supplemented with 10% FBS and 1% penicillin/streptomycin at 37 °C on a heated stage in an environmental chamber (TOKAI HIT Co., Ltd.). Images were acquired with a microscope (ECLIPSE Ti2) with a 100x 1.4 NA objective, an 16 XY Piezo-Z stage (Nikon instruments Inc.), a spinning disk (Yokogawa), an electron multiplier charge-coupled device camera (IXON-L-897) and a laser merge module equipped with 488nm, 561nm, 594 nm, and 630nm lasers controlled by NIS-Elements Advanced Research. For fixed cells, images were taken with 0.5 uM spacing between Z slices, for a total of 8 uM. For single-particle tracking, GFP images were taken at two time intervals, 30ms and 5s, to generate tracks for 0-1s timescale and 10-100 s timescale. For 5 s time interval, both GFP and mCherry images were taken during the time course to accurately identify TERRA telomeric localization. For 30 ms interval, taking a mCherry image before and after tracking the GFP channel was sufficient to determine co-localization between TERRA foci and telomeres.

### Chemical dimerization

Dimerization on telomeres was performed as previously described (Zhao et al., 2021). Briefly, dimerizers were added to growth medium to a final working concentration of 100 nM in a dark room with a dim red-light lamp. Cells were incubated with the dimerizers-containing medium for 2 h, followed by immunofluorescence (IF) or fluorescence in situ hybridization (FISH).

### Image processing

Images were processed and analyzed using NIS Elements Software. Maximum projections were created from z stacks, and thresholds were applied to the resulting 2D images to segment and identify TERRA foci as binaries. For colocalization quantification of two fluorescent labels, fixed images were analyzed by NIS-Elements AR to determine if the different labels were located in the same area of the cell.

### Single-particle tracking

NIS Elements tracking module was used to generate tracks for the TERRA binaries and Mean Square Displacements (MSDs) were calculated as:

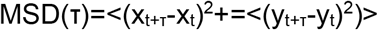

Where x_t_ and y_t_ are the foci coordinates at time t while x_t+τ_ and y_t+τ_ are the foci coordinates after a lag time of *τ*. MATLAB was used for plotting and curve-fitting MSD vs. Lag Time log-log plots to evaluate the anomalous exponent α from the anomalous diffusion model:

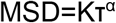

Where K is the generalized diffusion coefficients, *τ* is the lag time, and α is the anomalous exponent. The time-dependent diffusion coefficient, D, is calculated from mean MSD at each *τ* from MSD=4D*τ*.

### Statistical analyses

All p values were generated with a two-sample t-test in MATLAB with function ttest2.

## Acknowledgment

We thank Dr. Bruce Armitage for helpful discussions and Dr. Tumul Srivastava for making the TERRA FISH probe. We thank Dr. Lingling Chen for kindly providing us with the protocol for dynamic RNA imaging in living cells with CRISPR-Cas13 system. This work is supported by the US National Institutes of Health (U01CA260851 to H.Z., GM118510 to D.M.C.).

---

**Supplemental figure S1.**
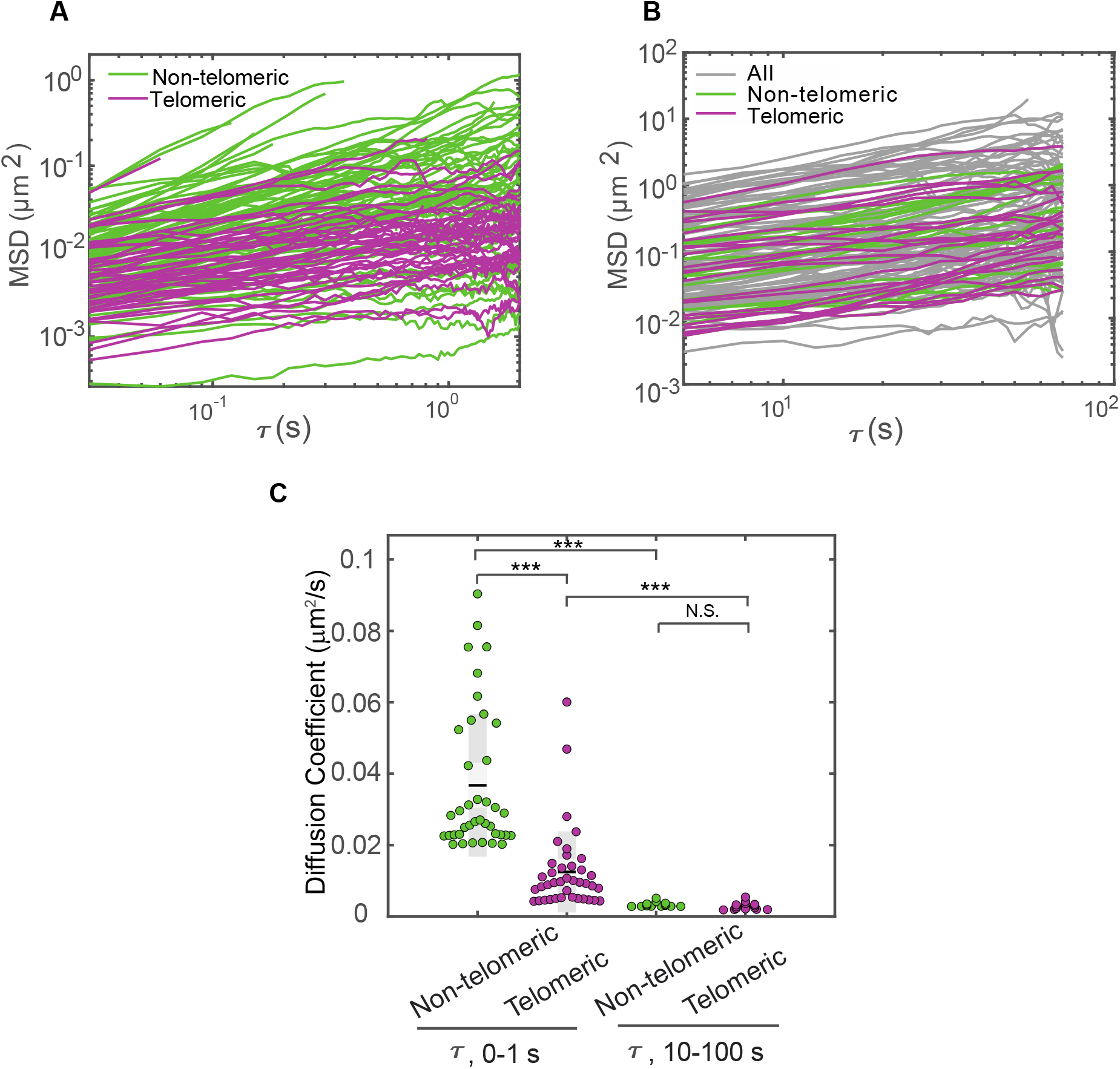
TERRA anomalous diffusion exponents and effective diffusion coefficients. (A) MSDs for non-telomeric and telomeric TERRA foci at timescale 0-1 s. (B) MSDs for non-telomeric and telomeric TERRA foci at timescale 10-100 s. (C) Average diffusion coefficients of non-telomeric and telomeric foci at timescale 0-1 s and 10-100 s. *** indicates p<0.001. N.S., not significant.

**Supplemental Table 1.** Anomalous exponent and mean diffusion coefficient in Figure 3D and Figure S1C.

|  | Time scale: 0-1 s |  | Time scale: 10-100 s |  |
| --- | --- | --- | --- | --- |
|  | Non-Telomeric | Telomeric | Non-Telomeric | Telomeric |
| Mean Anomalous Exponent ($\alpha$) | $0.52 \pm 0.04$ | $0.34 \pm 0.19$ | $0.70 \pm 0.33$ | $0.64 \pm 0.33$ |
| Mean Diffusion Coefficient ($\mu\text{m}^2/\text{s}$) | $0.037 \pm 0.020$ | $0.013 \pm 0.011$ | $0.0033 \pm 0.0007$ | $0.0029 \pm 0.001$ |

## Notes

### Competing Interest Statement

The authors have declared no competing interest.

## Reference

Arora, R., & Azzalin, C. M. (2015). Telomere elongation chooses TERRA ALTernatives. RNA Biology, 12(9), 938–941. https://doi.org/10.1080/15476286.2015.1065374

Avogaro, L., Querido, E., Dalachi, M., Jantsch, M. F., Chartrand, P., & Cusanelli, E. (2018). Live-cell imaging reveals the dynamics and function of single-telomere TERRA molecules in cancer cells. RNA Biology, 15(6), 787–796. https://doi.org/10.1080/15476286.2018.1456300

Azzalin, C. M., & Lingner, J. (2015). Telomere functions grounding on TERRA firma. In Trends in Cell Biology (Vol. 25, Issue 1, pp. 29–36). https://doi.org/10.1016/j.tcb.2014.08.007

Azzalin, C. M., Reichenbach, P., Khoriauli, L., Giulotto, E., & Lingner, J. (2007). Telomeric repeat-containing RNA and RNA surveillance factors at mammalian chromosome ends. Science, 318(5851), 798–801. https://doi.org/10.1126/science.1147182

Ballister, E. R., Aonbangkhen, C., Mayo, A. M., Lampson, M. A., & Chenoweth, D. M. (2014). Localized light-induced protein dimerization in living cells using a photocaged dimerizer. Nature Communications, 5, 1–9. https://doi.org/10.1038/ncomms6475

Beishline, K., Vladimirova, O., Tutton, S., Wang, Z., Deng, Z., & Lieberman, P. M. (2017). CTCF driven TERRA transcription facilitates completion of telomere DNA replication. Nature Communications, 8(1), 1–10. https://doi.org/10.1038/s41467-017-02212-w

Bettin, N., Oss Pegorar, C., & Cusanelli, E. (2019). The Emerging Roles of TERRA in Telomere Maintenance and Genome Stability. Cells, 8(3), 246. https://doi.org/10.3390/cells8030246

Biffi, G., Tannahill, D., & Balasubramanian, S. (2012). An intramolecular G-quadruplex structure is required for binding of telomeric repeat-containing RNA to the telomeric protein TRF2. Journal of the American Chemical Society, 134(29), 11974–11976. https://doi.org/10.1021/ja305734x

Bonnell, E., Pasquier, E., & Wellinger, R. J. (2021). Telomere Replication: Solving Multiple End Replication Problems. Frontiers in Cell and Developmental Biology, 9(April), 1–17. https://doi.org/10.3389/fcell.2021.668171

Chu, H. P., Cifuentes-Rojas, C., Kesner, B., Aeby, E., Lee H. goo, Wei, C., Oh, H. J., Boukhali, M., Haas, W., & Lee, J. T. (2017). TERRA RNA Antagonizes ATRX and Protects Telomeres. Cell, 170(1), 86-101.e16. https://doi.org/10.1016/j.cell.2017.06.017

Claude, E., & Decottignies, A. (2020). Telomere maintenance mechanisms in cancer: telomerase, ALT or lack thereof. Current Opinion in Genetics and Development, 60, 1–8. https://doi.org/10.1016/j.gde.2020.01.002

De Silanes, I. L., D’Alcontres, M. S., & Blasco, M. A. (2010). TERRA transcripts are bound by a complex array of RNA-binding proteins. Nature Communications, 1(3), 1–10. https://doi.org/10.1038/ncomms1032

De Silanes, I. L., Graña, O., De Bonis, M. L., Dominguez, O., Pisano, D. G., & Blasco, M. A. (2014). Identification of TERRA locus unveils a telomere protection role through association to nearly all chromosomes. Nature Communications, 5, 1–13. https://doi.org/10.1038/ncomms5723

Deng, Z., Norseen, J., Wiedmer, A., Riethman, H., & Lieberman, P. M. (2009). TERRA RNA binding to TRF2 facilitates heterochromatin formation and ORC recruitment at telomeres. Mol Cell, 35(4), 403–413. https://doi.org/10.1016/j.molcel.2009.06.025

Dilley, R. L., & Greenberg, R. A. (2015). ALTernative Telomere Maintenance and Cancer. Trends in Cancer, 20(2), 163–178. https://doi.org/10.1016/j.trecan.2015.07.007.ALTernative

Diman, A., & Decottignies, A. (2018). Genomic origin and nuclear localization of TERRA telomeric repeat-containing RNA: from Darkness to Dawn. FEBS Journal, 285(8), 1389–1398. https://doi.org/10.1111/febs.14363

Feretzaki, M., Pospisilova, M., Valador Fernandes, R., Lunardi, T., Krejci, L., & Lingner, J. (2020). RAD51-dependent recruitment of TERRA lncRNA to telomeres through R-loops. Nature, 587, 303–308. https://doi.org/10.1038/s41586-020-2815-6

Flynn, R. L., Centore, R. C., O’Sullivan, R. J., Rai, R., Tse, A., Songyang, Z., Chang, S., Karlseder, J., & Zou, L. (2011). TERRA and hnRNPA1 orchestrate an RPA-to-POT1 switch on telomeric single-stranded DNA. Nature, 471(7339), 532–538. https://doi.org/10.1038/nature09772

Frank, L., & Rippe, K. (2020). Repetitive RNAs as Regulators of Chromatin-Associated Subcompartment Formation by Phase Separation. Journal of Molecular Biology, 432(15), 4270–4286. https://doi.org/10.1016/j.jmb.2020.04.015

Hanahan, D., & Weinberg, R. A. (2011). Hallmarks of cancer: The next generation. Cell, 144(5), 646–674. https://doi.org/10.1016/j.cell.2011.02.013

Jain, A., & Vale, R. D. (2017). RNA phase transitions in repeat expansion disorders. Nature,546, 243–247. https://doi.org/10.1038/nature22386

Jegou, T., Chung, I., Heuvelman, G., Wachsmuth, M., Gorisch, S. M., Greulich-Bode, K. M., Boukamp, P., Lichter, P., & Rippe, K. (2009). Dynamics of Telomeres and Promyelocytic Leukemia Nuclear Bodies in a Telomerase-negative Human Cell Line. Molecular Biology of the Cell, 20, 2070–2082. https://doi/pdf/10.1091/mbc.e08-02-0108

Khandelia, P., Yap, K., & Makeyev, E. V. (2011). Streamlined platform for short hairpin RNA interference and transgenesis in cultured mammalian cells. Proceedings of the National Academy of Sciences of the United States of America, 108(31), 12799–12804. https://doi.org/10.1073/pnas.1103532108

Lalonde, M., & Chartrand, P. (2020). TERRA, a Multifaceted Regulator of Telomerase Activity at Telomeres. Journal of Molecular Biology, 432(15), 4232–4243. https://doi.org/10.1016/j.jmb.2020.02.004

Maciejowski, J., & De Lange, T. (2017). Telomeres in cancer: Tumour suppression and genome instability. Nature Reviews Molecular Cell Biology, 18(3), 175–186. https://doi.org/10.1038/nrm.2016.171

Marión, R. M., Montero, J. J., de Silanes, I. L., Graña-Castro, O., Martínez, P., Schoeftner, S., Palacios-Fábrega, J. A., & Blasco, M. A. (2019). TERRA regulate the transcriptional landscape of pluripotent cells through TRF1-dependent recruitment of PRC2. ELife, 8, 1–32. https://doi.org/10.7554/eLife.44656

Mei, Y., Deng, Z., Vladimirova, O., Gulve, N., Johnson, F. B., Drosopoulos, W. C., Schildkraut, C. L., & Lieberman, P. M. (2021). TERRA G-quadruplex RNA interaction with TRF2 GAR domain is required for telomere integrity. Scientific Reports, 11(1), 1–14. https://doi.org/10.1038/s41598-021-82406-x

Min, J., Wright, W. E., & Shay, J. W. (2019). Clustered telomeres in phase-separated nuclear condensates engage mitotic DNA synthesis through BLM and RAD52. Genes and Development, 33(13–14), 814–827. https://doi.org/10.1101/gad.324905.119

Montero, J. J., López-Silanes, I., Megías, D., Fraga, M., Castells-García, Á., & Blasco, M. A. (2018). TERRA recruitment of polycomb to telomeres is essential for histone trymethylation marks at telomeric heterochromatin. Nature Communications, 1548(9). https://doi.org/10.1038/s41467-018-03916-3

O’Sullivan, R. J., & Karlseder, J. (2010). Telomeres: Protecting chromosomes against genome instability. Nature Reviews Molecular Cell Biology, 11(3), 171–181. https://doi.org/10.1038/nrm2848

Palm, W., & de Lange, T. (2008). How Shelterin Protects Mammalian Telomeres. Annual Review of Genetics, 42(1), 301–334. https://doi.org/10.1146/annurev.genet.41.110306.130350

Pédelacq, J. D., Cabantous, S., Tran, T., Terwilliger, T. C., & Waldo, G. S. (2006). Engineering and characterization of a superfolder green fluorescent protein. Nature Biotechnology, 24(1), 79–88. https://doi.org/10.1038/nbt1172

Petti, E., Buemi, V., Zappone, A., Schillaci, O., Broccia, P. V., Dinami, R., Matteoni, S., Benetti, R., & Schoeftner, S. (2019). SFPQ and NONO suppress RNA:DNA-hybrid-related telomere instability. Nature Communications, 10(1). https://doi.org/10.1038/s41467-019-08863-1

Porreca, R. M., Herrera-Moyano, E., Skourti, E., Law, P. P., Franco, R. G., Montoya, A., Faull, P., Kramer, H., & Vannier, J. B. (2020). Trf1 averts chromatin remodelling, recombination and replication dependent-break induced replication at mouse telomeres. ELife, 9, 1–28. https://doi.org/10.7554/eLife.49817

Recagni, M., Bidzinska, J., Zaffaroni, N., & Folini, M. (2020). The role of alternative lengthening of telomeres mechanism in cancer: Translational and therapeutic implications. Cancers, 12(4), 1–15. https://doi.org/10.3390/cancers12040949

Schoeftner, S., & Blasco, M. A. (2008). Developmentally regulated transcription of mammalian telomeres by DNA-dependent RNA polymerase II. Nature Cell Biology, 10(2), 228–236. https://doi.org/10.1038/ncb1685

Silva, B., Arora, R., Bione, S., & Azzalin, C. M. (2021). TERRA transcription destabilizes telomere integrity to initiate break-induced replication in human ALT cells. Nature Communications, 12(1), 1–12. https://doi.org/10.1038/s41467-021-24097-6

Soranno, A., Incicco, J. J., Bona P. De, Tomko, E. J., Galburt, E. A., Alex, S., & Galletto, R. (2021). Shelterin components modulate nucleic acids condensation and phase separation. BioRxiv Preprint, 18, 1–21. https://doi.org/10.1101/2021.04.30.442189

Tanenbaum, M. E., Gilbert, L. A., Qi, L. S., Weissman, J. S., & Vale, R. D. (2014). A protein-tagging system for signal amplification in gene expression and fluorescence imaging. Cell, 159(3), 635–646. https://doi.org/10.1016/j.cell.2014.09.039

Tseng, Y., Lee, J. S. H., Kole, T. P., Jiang, I., & Wirtz, D. (2004). Micro-organization and visco-elasticity of the interphase nucleus revealed by particle nanotracking. Journal of Cell Science, 117(10), 2159–2167. https://doi.org/10.1242/jcs.01073

Wang, C., Zhao, L., & Lu, S. (2015). Role of TERRA in the regulation of telomere length. International Journal of Biological Sciences, 11(3), 316–323. https://doi.org/10.7150/ijbs.10528

Woringer, M., & Darzacq, X. (2018). Protein motion in the nucleus: from anomalous diffusion to weak interactions. Biochemical Society Transactions, 46(4), 945–956. https://doi.org/10.1042/BST20170310

Yamada, T., Yoshimura, H., Shimada, R., Hattori, M., Eguchi, M., Fujiwara, T. K., Kusumi, A., & Ozawa, T. (2016). Spatiotemporal analysis with a genetically encoded fluorescent RNA probe reveals TERRA function around telomeres. Scientific Reports, 6(August), 1–13. https://doi.org/10.1038/srep38910

Yang, L. Z., Wang, Y., Li, S. Q., Yao, R. W., Luan, P. F., Wu, H., Carmichael, G. G., & Chen, L. L. (2019). Dynamic Imaging of RNA in Living Cells by CRISPR-Cas13 Systems. Molecular Cell, 76(6), 981-997.e7. https://doi.org/10.1016/j.molcel.2019.10.024

Yeager, T. R., Neumann, A. A., Englezou, A., Huschtscha, L. I., Noble, J. R., & Reddel, R. R. (1999). Telomerase-negative immortalized human cells contain a novel type of promyelocytic leukemia (PML) body. Cancer Research, 59(17), 4175–4179.

Yu, J. (2016). Single-Molecule Studies in Live Cells. Annual Review of Physical Chemistry, 67, 565–585. https://doi.org/10.1146/annurev-physchem-040215-112451

Zhang, H., Aonbangkhen, C., Tarasovetc, E. V., Ballister, E. R., Chenoweth, D. M., & Lampson, M. A. (2017). Optogenetic control of kinetochore function. Nature Chemical Biology, 13(10), 1096–1101. https://doi.org/10.1038/nchembio.2456

Zhang, H., Elbaum-Garfinkle, S., Langdon, E. M., Taylor, N., Occhipinti, P., Bridges, A. A., Brangwynne, C. P., & Gladfelter, A. S. (2015). RNA Controls PolyQ Protein Phase Transitions. Molecular Cell, 60(2), 220–230. https://doi.org/10.1016/j.molcel.2015.09.017

Zhang, H., Zhao, R., Tones, J., Liu, M., Dilley, R. L., Chenoweth, D. M., Greenberg, R. A., & Lampson, M. A. (2020). Nuclear body phase separation drives telomere clustering in ALT cancer cells. Molecular Biology of the Cell, 31(18), 2048–2056. https://doi.org/10.1091/mbc.E19-10-0589

Zhang, J. M., Yadav, T., Ouyang, J., Lan, L., & Zou, L. (2019). Alternative Lengthening of Telomeres through Two Distinct Break-Induced Replication Pathways. Cell Reports, 26(4), 955-968.e3. https://doi.org/10.1016/j.celrep.2018.12.102

Zhao, R., Chenoweth, D. M., & Zhang, H. (2021). Chemical dimerization-induced protein condensates on telomeres. Journal of Visualized Experiments, 2021(170). https://doi.org/10.3791/62173

